# Longitudinal alterations in fronto-striatal glutamate are associated with functioning during inhibitory control in autism spectrum disorder and obsessive compulsive disorder

**DOI:** 10.1101/2020.04.14.031740

**Authors:** Viola Hollestein, Jan K. Buitelaar, Daniel Brandeis, Tobias Banaschewski, Anna Kaiser, Sarah Hohmann, Bob Oranje, Bram Gooskens, Sarah Durston, David J. Lythgoe, Jilly Naaijen

## Abstract

**Background:** Autism spectrum disorder (ASD) and obsessive compulsive disorder (OCD) are neurodevelopmental disorders with overlapping symptomatology. Both show deficits in inhibitory control, which are associated with altered functioning and glutamate concentrations in the fronto-striatal circuitry. These parameters have never been examined together. Here we, for the first time, used a multi-center, longitudinal approach to investigate fronto-striatal functioning during an inhibitory control task and its association with fronto-striatal glutamate concentrations across these two disorders.

**Methods:** 74 adolescents with ASD (24) or OCD (15) and controls (35) aged 8-17 were recruited across three sites of the European TACTICS consortium. They underwent two magnetic resonance imaging (MRI) sessions with a one-year interval. This included proton magnetic resonance spectroscopy (^1^H-MRS; n=74) and functional MRI during an inhibitory control task (n=57). We used linear mixed effects models to investigate, over time, the relationship between fronto-striatal functioning and glutamate concentrations across these groups and continuous measures of overlapping compulsivity symptoms.

**Results:** During failed inhibitory control, in OCD increased striatal glutamate was associated with increased neural activation of ACC, an effect that decreased over time. During successful inhibitory control, higher ACC glutamate was positively associated with striatal activation in OCD and compulsivity across time. ACC glutamate levels decreased over time in the ASD group compared to controls, while striatal glutamate decreased over time, independent of diagnosis.

**Conclusions:** Significant differences in fronto-striatal glutamate were observed in ASD and OCD, affecting functional activity during failed- and successful inhibitory control differently, especially in OCD, with effects changing over time.

## Introduction

Although autism spectrum disorder (ASD) and obsessive compulsive disorder (OCD) are two separate neurodevelopmental disorders with distinct diagnostic characteristics (1), they are highly comorbid and a comparison of symptoms has suggested that there are more similarities than differences between them (2–4). Individuals with these disorders typically show compulsive behaviors, which are defined as the repetitive, irresistible urge to perform certain behaviors or thoughts, and diminished control over this urge (5). Compulsive behaviors are associated with deficits in inhibitory control in tasks such as the stop-signal task (3, 6). Fronto-striatal areas are known to be involved in inhibitory control, and are strongly regulated by the neurotransmitter glutamate (7–9). Structural, functional and neurochemical imaging studies show alterations across fronto-striatal brain regions in both ASD and OCD, which suggest a possible shared underlying mechanism affecting compulsive behaviors (10–12).

Compulsive behaviors are highly associated with deficits in inhibitory control, which in turn is regulated by the fronto-striatal circuitry, where particularly the anterior cingulate cortex (ACC) is crucial for exerting inhibitory control, and the striatum is thought to be important driving them (7, 10, 12–16).

In studies using the stop signal task in ASD and OCD, there has been inconsistent results. Some studies have found no behavioral differences in ASD nor in OCD (17–19), while others have found worse performance in participants with OCD (5, 6, 20–22), demonstrating deficits in inhibitory control. However, these differences are more commonly found in adults with OCD than children (23). Altered activity in fronto-striatal areas during response inhibition has been found in both disorders as well (18, 24, 25) and some studies found altered functional activity despite not finding behavioral differences in response inhibition compared to controls (26, 27). However, in a previous study using a partly overlapping sample of the current study, no behavioral or neural alterations were found during inhibitory control in participants with ASD and OCD (19).

Several studies suggest that altered glutamate concentrations in the fronto-striatal regions may be associated with deficits in inhibitory control. For instance, altered glutamate concentrations have been linked to repetitive behaviors and compulsivity (7, 28), which seem to differ in individuals with ASD and OCD compared to controls across development. A meta-analysis of studies investigating fronto-striatal glutamate using proton magnetic resonance spectroscopy (^1^H-MRS) in neurodevelopmental disorders reported that increased glutamate concentrations in striatum seemed to be present across both disorders (7). In the ACC, on the other hand, glutamate concentrations were often higher in children and adolescents with these disorders while in adults the opposite pattern was found, with lower concentrations compared to controls, suggesting a developmental shift (7).

In a study investigating glutamate concentrations and neural functioning during inhibitory control in an ADHD population, decreased ACC glutamate was associated with increased activity in the striatum (9).

This strongly suggests that investigating the interplay between glutamate and functional activity during inhibitory control is an important step for understanding the mechanistic underpinnings of behaviors across neurodevelopmental disorders. In a study including the first time of measure (T1) of the participants in this study, increased ACC glutamate was found in both ASD and OCD, and a positive association between ACC glutamate and compulsivity was found (8). In the current study we followed part of this multi-center, multimodal sample up (T2). With this longitudinal data we aim to investigate whether fronto-striatal glutamatergic alterations and functioning during inhibitory control are stable across (atypical) neurodevelopment. In addition, we were interested in the relation between these parameters and expected fronto-striatal glutamatergic alterations to be negatively associated with neural activation during inhibitory control across ASD and OCD.

## Methods and Materials

### Participants

We included 74 participants (ASD = 24, OCD = 15, controls = 35) for the longitudinal ^1^H-MRS analysis, who were between 8 and 16 years old at the first time of measurement, and between 9 and 17 years at the second measurement. The paper regarding T1 of our sample included a total amount of n=133 participants (Naaijen et al., 2017). Reasons for drop-out for this longitudinal study were loss of interest or quality issues regarding ^1^H-MRS (Nijmegen = 19, Mannheim = 11, London = 13, Utrecht = 33). For the combined ^1^H-MRS and fMRI analysis we included 57 participants (for more details see the next section). The participants were recruited at three different locations across Europe (Radboud University Medical Center and the Donders Institute for Brain, Cognition and Behavior, Nijmegen, The Netherlands (N = 38), Kings College London, London, United Kingdom (N = 17), Central Institute of Mental Health, Mannheim, Germany (N = 19)) in the multicenter study COMPULS, part of the TACTICS consortium (www.tactics-project.eu). Another site was excluded due to too few participants surviving quality control (N=3). The inclusion criteria were IQ > 70, ability to speak and comprehend the native language of the location of recruitment and being of Caucasian descent (for further details, see (29)). To confirm DSM-IV-TR (30) diagnoses of ASD and OCD, we used the Autism Diagnostic Interview-Revised (ADI-R) (31) (and Children’s Yale Brown Obsessive Compulsive Scale (CYBOCS) (32) for ASD and OCD respectively. Control participants were confirmed to not score in the clinical range for any diagnoses using the Child Behavior Checklist (CBCL) and the Teacher Report Form (TRF) (33). Compulsive behaviors were measured using the Repetitive Behavior Scale – Revised (RBS-R) (34). Information on medication use was collected on the measurement days via parental report. Participants were asked to abstain from stimulant medication 48 hours before scanning. Ethical approval for the study was obtained for all centers separately and participants and their parents gave written informed consent for participation.

### Stop-Signal Task

To measure inhibitory control participants completed a modified visual version of the stop-signal task (SST) (35) during an fMRI session. For details of the design of the task, see Figure 1. For details of behavioral measures and results of the behavioral analysis, see the supplementary material.

**Figure 1:**
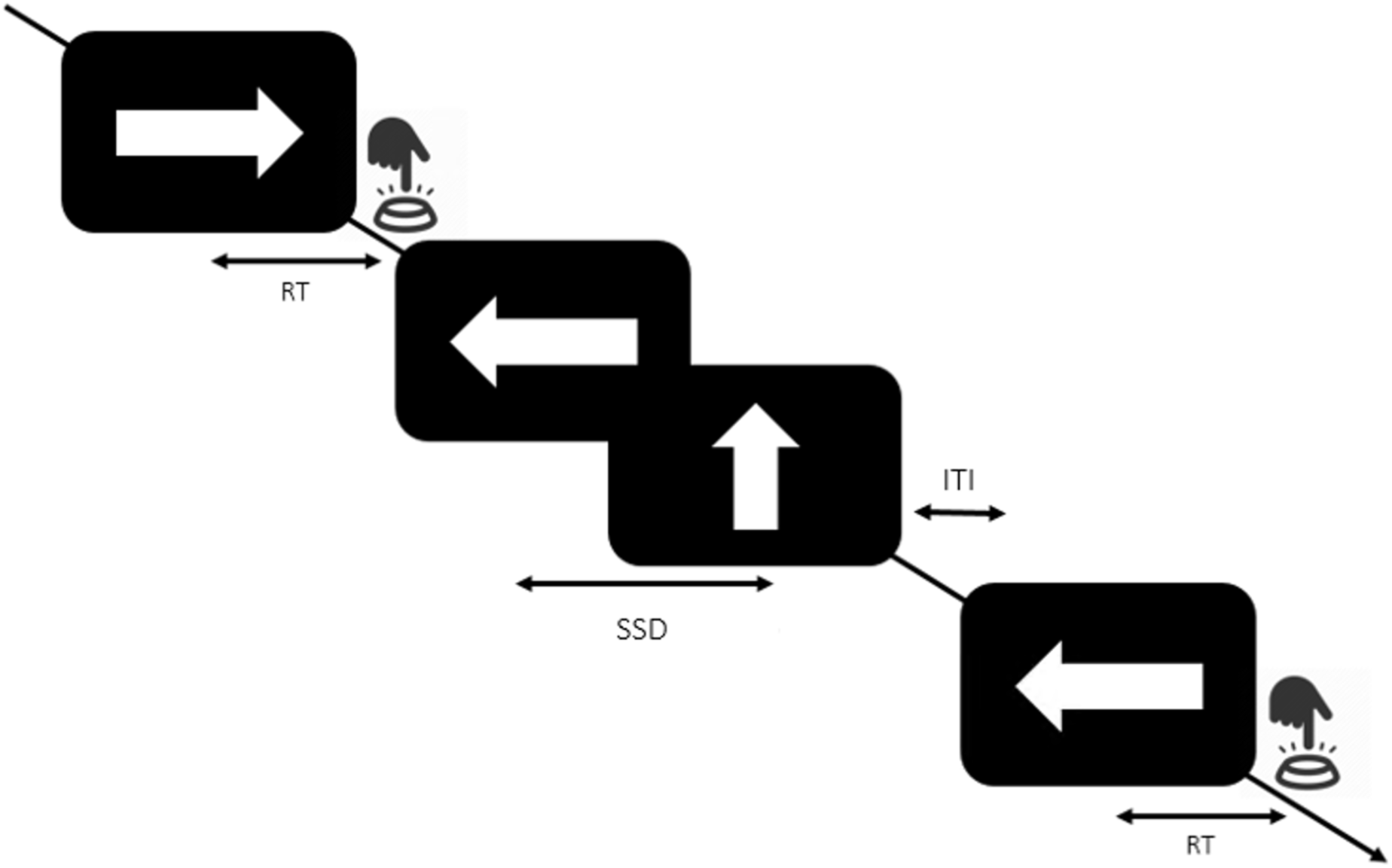
Stop-signal task. Arrows were presented on a screen; the task was to press a button indicating the direction the arrow was pointing at. In 20% of trials the arrow was followed by a stop cue of an arrow pointing upwards, instructing to withhold a response. The stop-signal delay (SSD) between stimulus onset and stop-signal was adaptive, where the SSD after successful inhibition increased with 50 ms while after failed inhibition it decreased with 50 ms. This ensured participants success to inhibit in approximately 50% of the stop-trials. The inter-trial interval (ITI), the time between the trials, was randomly jittered between 1.6 and 2.0 seconds.

### Image Acquisition

Participants were familiarized with the MRI settings and practice of the SST using a dummy scanner at T1. The data were acquired from the three study locations, all using 3 Tesla scanners (Siemens Trio and Siemens Prisma, Siemens, Erlangen, Germany; Philips 3 T Achieva, Philips Medical Systems, Best, The Netherlands; General Electric MR750, GE Medical Systems, Milwaukee, Wi, USA). Structural T1-weighted scans were acquired based on the ADNI GO protocols (36, 37), which were used for registration of the functional scans and voxel placement for the ^1^H-MRS. Details on the structural, functional and ^1^H-MRS scan parameters can be found in Table 1.

**Table 1:** Scan sequences.

| Sequence | Site | TR/TE/TI<br>(ms) | Flip<br>angle | Field<br>of<br>view<br>(mm) | Matrix<br>RL/AP/slice<br>s | Voxel –<br>size (mm) | Gap<br>(%) | Parallel<br>Imagin<br>g | Averages<br>Water<br>suppressed/<br>unsuppressed |
| --- | --- | --- | --- | --- | --- | --- | --- | --- | --- |
|  |  |  |  |  |  |  |  |  | d |
| T1 | Nijmegen | 2300*/2.98/90 | 9 | 256 | 212/256/176 | 1.0*1.0*1. | NA | 2 | NA |
|  | (Siemens) | 0 |  |  |  | 2 |  |  |  |
|  | Mannhei | 2300*/2.98/90 | 9 | 270 | 212/254/176 | 1.1*1.1*1. | NA | 2 | NA |
|  | m | 0 |  |  |  | 2 |  |  |  |
|  | (Siemens) |  |  |  |  |  |  |  |  |
|  | London | 7.31*/3.02/400 | 11 | 270 | 256/256/196 | 1.1*1.1*1. | NA | 1.75 | NA |
|  | (GE) |  |  |  |  | 2 |  |  |  |
| MRS<br>PRESS | All | 3000/30/- | NA | NA | NA | 20*20*20 | NA | NA | 96/16 |
| Functiona<br>l MRI | All | 2070/35/- | 74 | 192 | 192/192/36 | 3.0*3.0*3. | 13 | 2 | NA |
|  |  |  |  |  |  | 0 |  |  |  |

\*As provided by the manufacturer. GE defines a TR as the time between excitation pulses, while Siemens defines TR as the time between inversion recovery pulses.

Glutamate concentrations were measured using ^1^H-MRS at the midline pregenual ACC and left dorsal striatum covering caudate and putamen, with a voxel size of 8cm^3^ (2’2’2). Voxel locations were adjusted to maximize the grey matter (GM) content and minimize the cerebrospinal fluid (CSF) content to keep the quality of the data as high as possible. The locations of all voxel placements are shown in Figure 2. Across all sites, the proton spectra were acquired using a point resolved spectroscopy sequence (PRESS) with the chemically selective suppression (CHESS) water suppression technique (38), details can be seen in Table 1.

**Figure 2:**
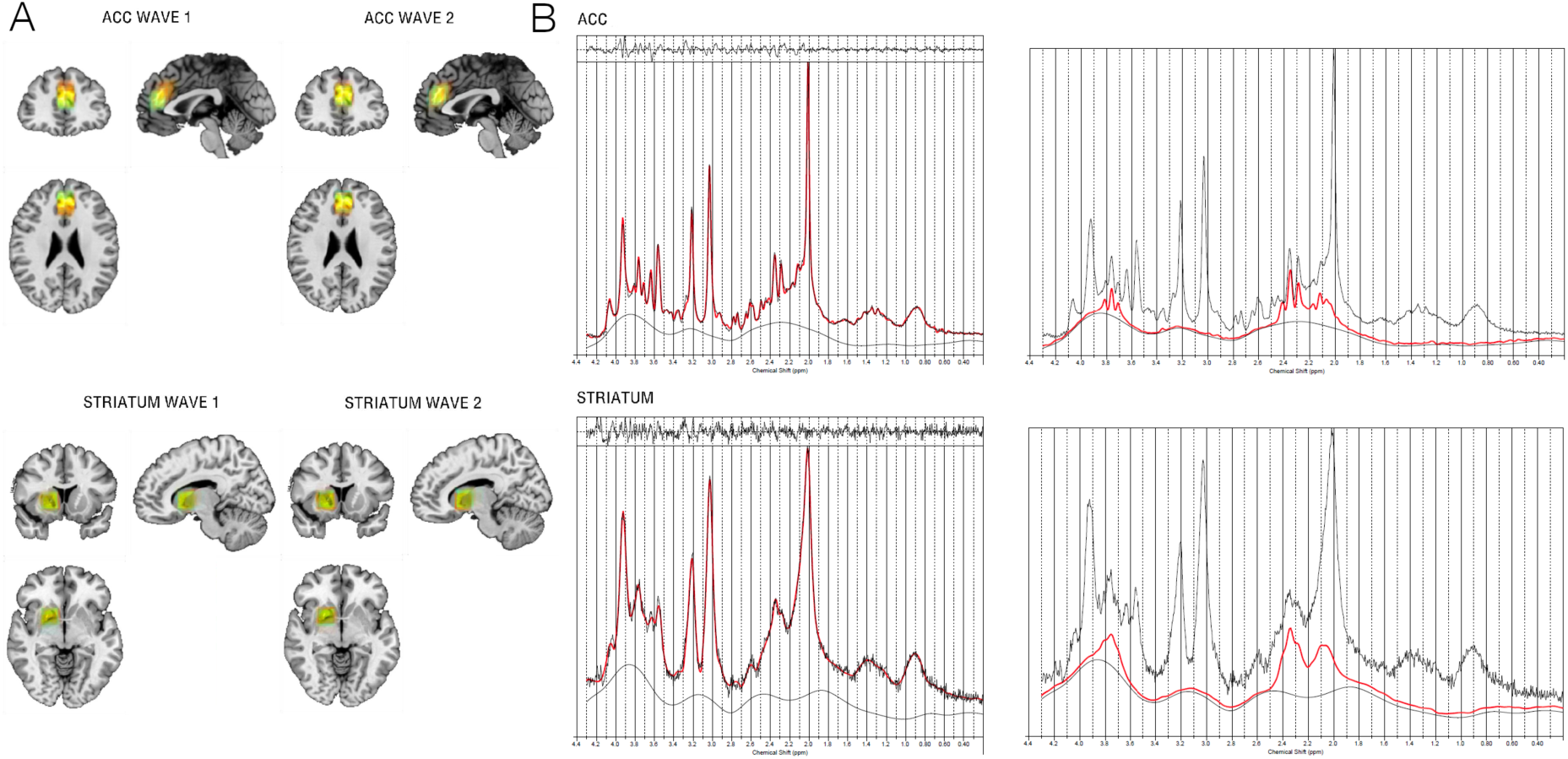
A: Superposition on the MNI152 template of all individual voxel placements in ACC and striatum, for ASD (red), OCD (blue) and controls (yellow). The placements were consistent across diagnoses, as seen by the large overlap of voxels. For voxel placements across sites, see Supplementary material. B: Example spectra of a 3T from proton magnetic resonance spectroscopy (^1^H-MRS) Linear Combination (LC) Model spectral fit in ACC and striatum from one of the control participants. The top of the images represents the residuals. The black line represents frequency-domain data, the red line is the LCModel fit. The right images show the fits for glutamate only. For examples of LCModel spectral fits and glutamate fits for each site, see Supplementary material.

### Imaging Analysis

#### fMRI

As per regular praxis, the first five volumes from each scan were removed to account for equilibration effects. Head movement correction was performed by realigning to the middle volume (MCFLIRT; (39)). A Gaussian kernel with full width at half maximum (FWHM) of 6 mm was used for grand mean scaling and spatial smoothing. ICA-AROMA (40, 41) was then used to remove signal components related to secondary-head motion artefacts, subsequently followed by nuisance regression to remove signal from CSF and white matter (WM), and high-pass filtering (100 sec). These images were then co-registered to each participants’ anatomical scan using boundary-based registration within FSL-FLIRT (42). The anatomical scans were spatially normalized using a 12-parameter affine registration to MNI152 standard space, using FSL-FNIRT (43). The images were then brought into standard space by applying the resulting warp fields to the concatenated functional image.

#### ^1^H-MRS

Glutamate concentrations were estimated using Linear Combination Model (LCModel) (44, 45). Example fitted spectra for both ACC and striatum can be seen in Figure 2. As different tissues contain different amounts of water, correction for tissue percentage and partial volume effects was calculated using the formula:

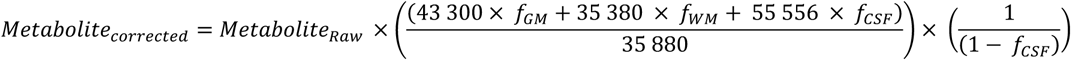

where 43 300 is the water concentration in millimolar for GM, 35 880 for WM and 55 556 for CSF, as described in the LCModel manual (44).

Participants were included when the signal-to-noise ratio was ≥ 15, Cramér-Rao lower bounds ≤ 20 %, or FWHM ≤ 0.1 parts per million. This resulted in 74 participants included in the analysis of ACC glutamate (ASD = 24, OCD = 15, controls = 35), and 56 participants included for striatal glutamate (Controls = 26, OCD = 12, ASD = 18).

### Statistical analyses

All analyses were performed using the R-software package (46) unless otherwise described.

We investigated fronto-striatal glutamate concentrations, neural activation and behavioral responses during inhibitory control separately. We tested whether glutamate concentrations (in either ACC or striatum) were associated with diagnosis, RBS-R total, RBS-R compulsivity subscale scores, time of measure (T1 and T2) and their possible interactions using linear mixed effects models (lme4 package (47)). Age, sex and site were added as covariates of non-interest and participant as a random factor to account for within subject variability across time. As age and sex did not affect the results or increased the fit of the model they were removed from further analyses.

Neural activation during inhibitory control was analyzed using SPM12 (Statistical Parametric Mapping release 12, https://www.fil.ion.ucl.ac.uk/spm/). For the whole brain analysis of fMRI during the stop-task, the first level models included two contrasts of interest; (1) failed stop – successful go, to isolate failed inhibitory control and (2) successful – failed stop, to isolate successful inhibitory control. For the second level of analysis looking at differences across groups and times of measure, t-contrasts were applied to these contrast maps. Covariates of non-interest were age, sex and site, to control for its possible effects

We additionally investigated whether the BOLD response in ACC and striatum during response inhibition was associated with glutamate concentrations in the same areas using diagnosis and time-point as additional predictors of interest. Here we used site as a covariate of non-interest and participant as the random factor as well. We used the average ^1^H-MRS voxels for ACC and striatum across participants as mapped onto the MNI152 space as our regions of interests (ROI) to extract the beta weights for our contrasts of interest. All reported *p*-values are corrected for multiple comparisons by the false discovery rate (FDR). Effect sizes are indicated as *r*, based on the function in the lme4 package (47).

## Results

### Demographics

No differences were found between the groups in age, IQ or sex. Both the RBS-R total score and all sub-scores were significantly different between controls and diagnostic groups where controls expectedly scored lower. There were no differences in the RBS total and sub-scores between T1 and T2. Table 2 shows an overview of the demographics and clinical variables of the largest subsample used.

**Table 2:**
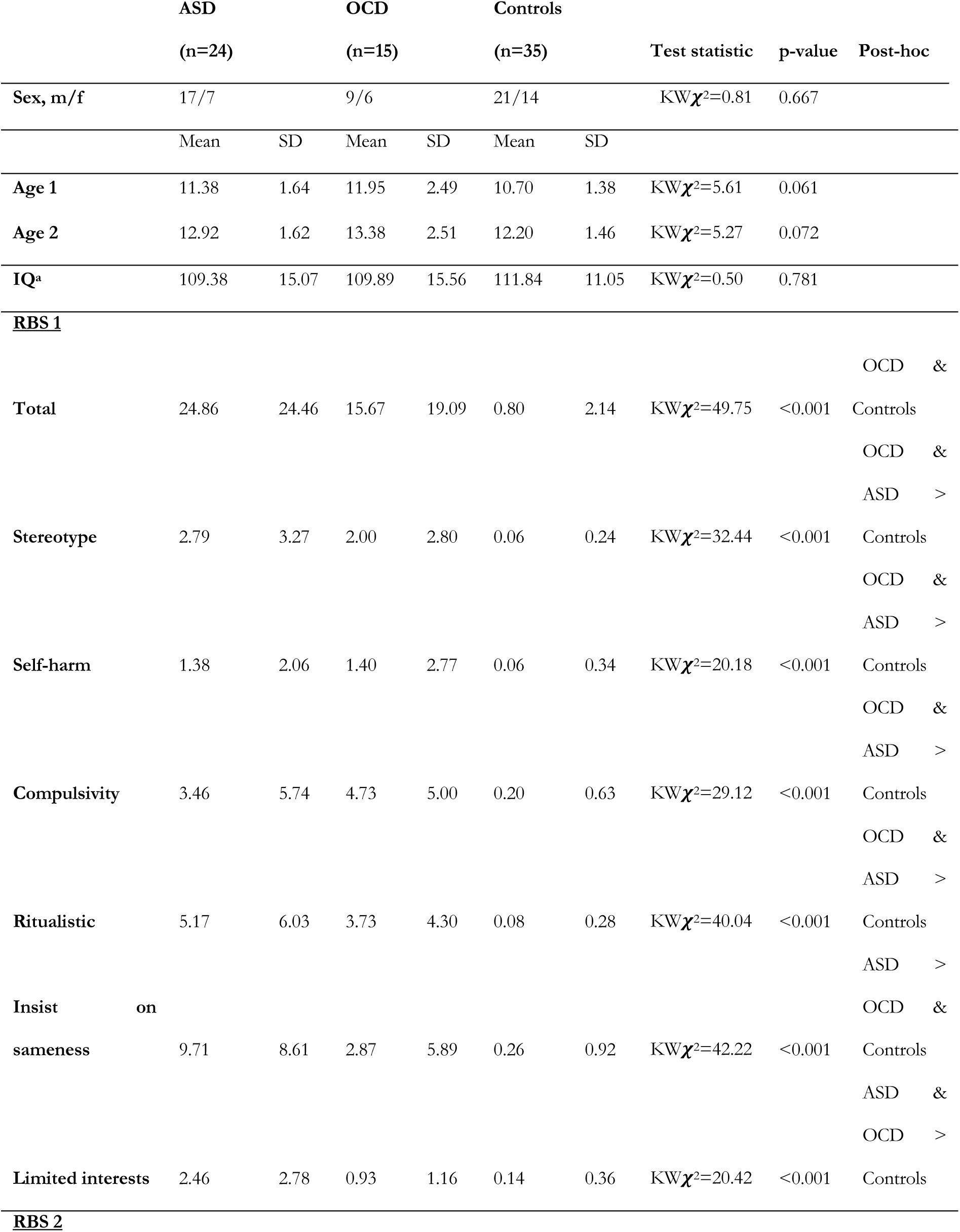
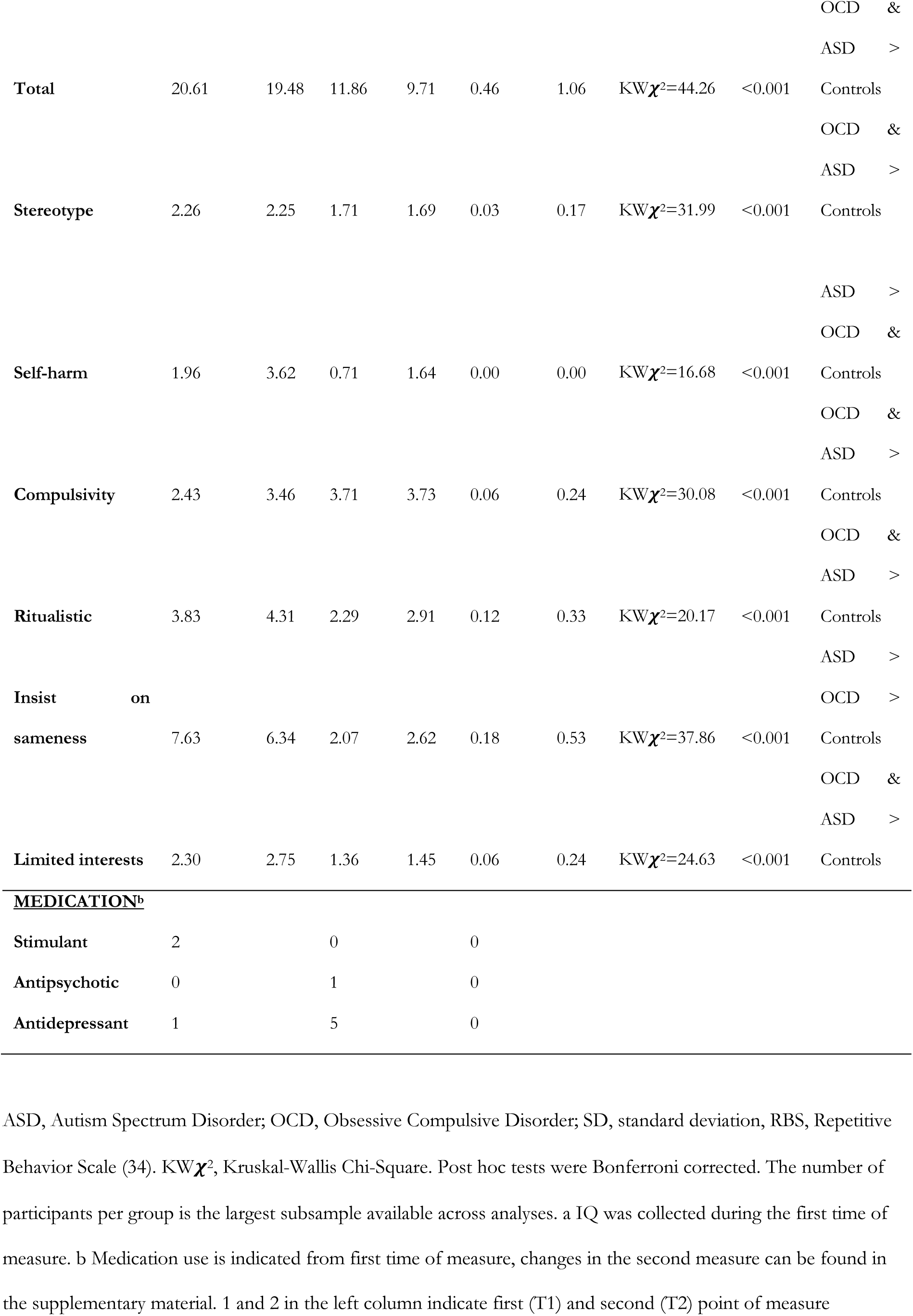
Demographic characteristics (based on the largest subsample group in analysis)

### Spectral quality

Groups did not differ in mean voxel percentage GM, WM or CSF in both voxels (all *p*-values > 0.05). Percentage GM in the striatum, however, was lower for the second time of measure compared to the first one ((b=-0.07, *t*_(52)_=- 2.97, *p* = 0.004), independent of diagnosis. To further verify that the spectral quality did not differ between groups, we compared the glutamate CRLB, overall signal to noise ratio (SNR) and linewidth (FWHM) of the spectra. No differences were found between the diagnostic groups or time-points for any of the measures. The ASD group showed, compared with controls, an increase in glutamate CRLB over time (b=0.009, *t*_(71)_=2.49, *p*=0.015), although with the highest CRLB of 14%, guaranteeing high quality of these spectra at both timepoints (48).

### Fronto-striatal glutamate

#### Group comparisons

There was an interaction between time and diagnosis associated with ACC glutamate. This interaction indicated a significantly larger decrease over time in ACC glutamate in ASD as compared to controls (b = −1.49, t_(71)_ = −2.26, *p* = 0.027, *r* = 0.26). No effects were found for the OCD group (all *p*-values > 0.05). No effect of diagnosis was found on striatal glutamate (all *p*-values > 0.05) but a significant overall effect of time was found (b = −0.65, t_(52)_ = −2.77, *p* = 0.016, *r* = 0.36) showing a decrease of glutamate over time (see Figure 3).

**Figure 3:**
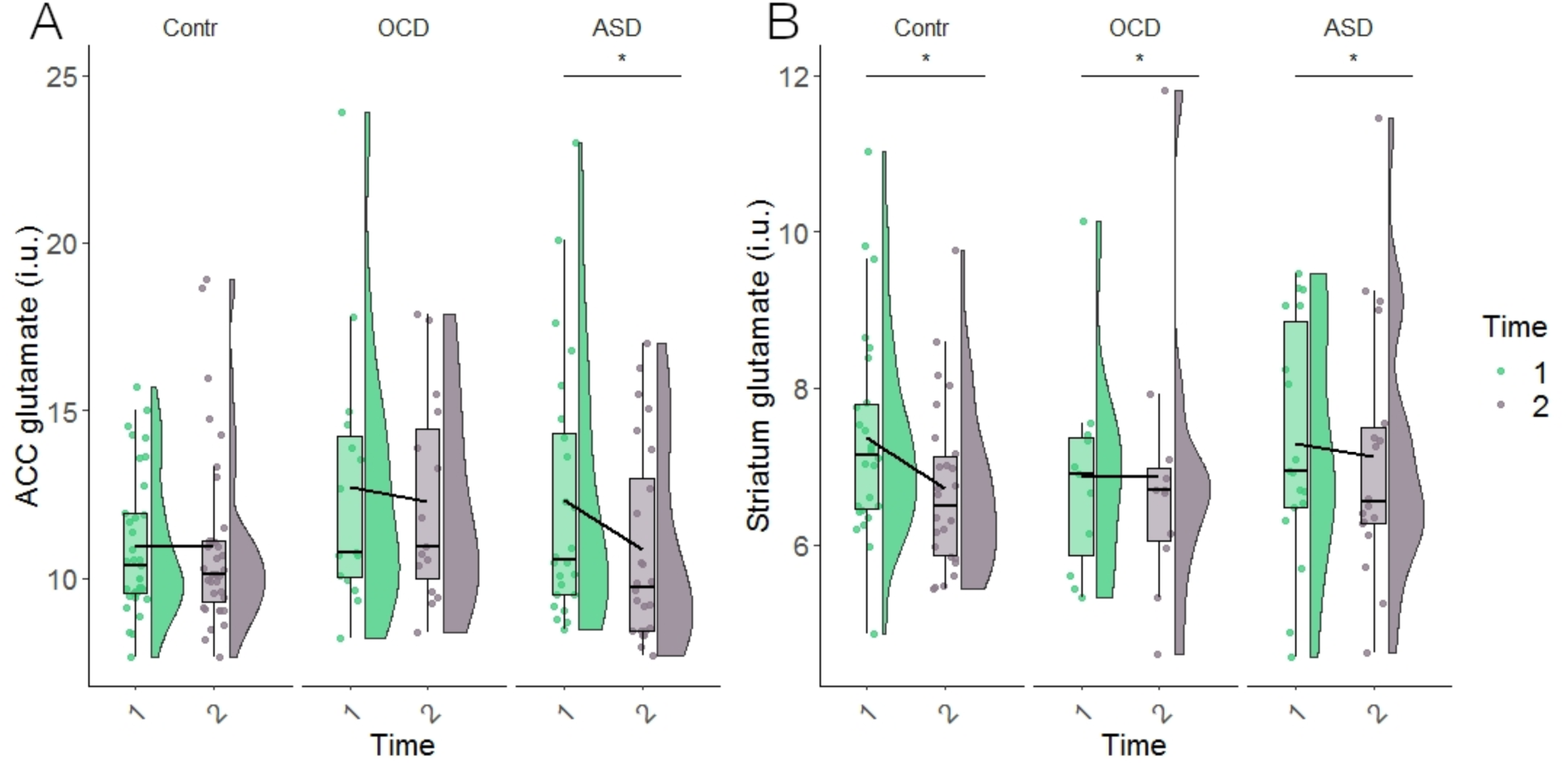
Glutamate concentrations in ACC (A) and striatum (B) across the groups. The asterisk in A indicate significantly larger difference in ACC glutamate over time in ASD compared to controls. The asterisks in B indicate significant difference in striatal glutamate over time independently of diagnosis. Plots were made using ggplot2 (54) and in-house adapted violin plots (55). *Note*: this figure shows raw data-points, not model estimates.

#### Continuous measures

No significant associations were found between the RBS total score, RBS compulsivity score and glutamate in ACC and striatum at either of the time-points separately nor longitudinally.

### Stop Signal Task

All groups showed common patterns of brain activation during successful inhibitory control and failed inhibitory control, where there was activation in areas typically associated with inhibitory control, such as ACC and striatum (Figure 4). No significant differences in neural activation between groups nor timepoints nor their interaction was found in our contrasts of interest (all *p*-values > 0.05). See the supplemental material for behavioral results.

**Figure 4:**
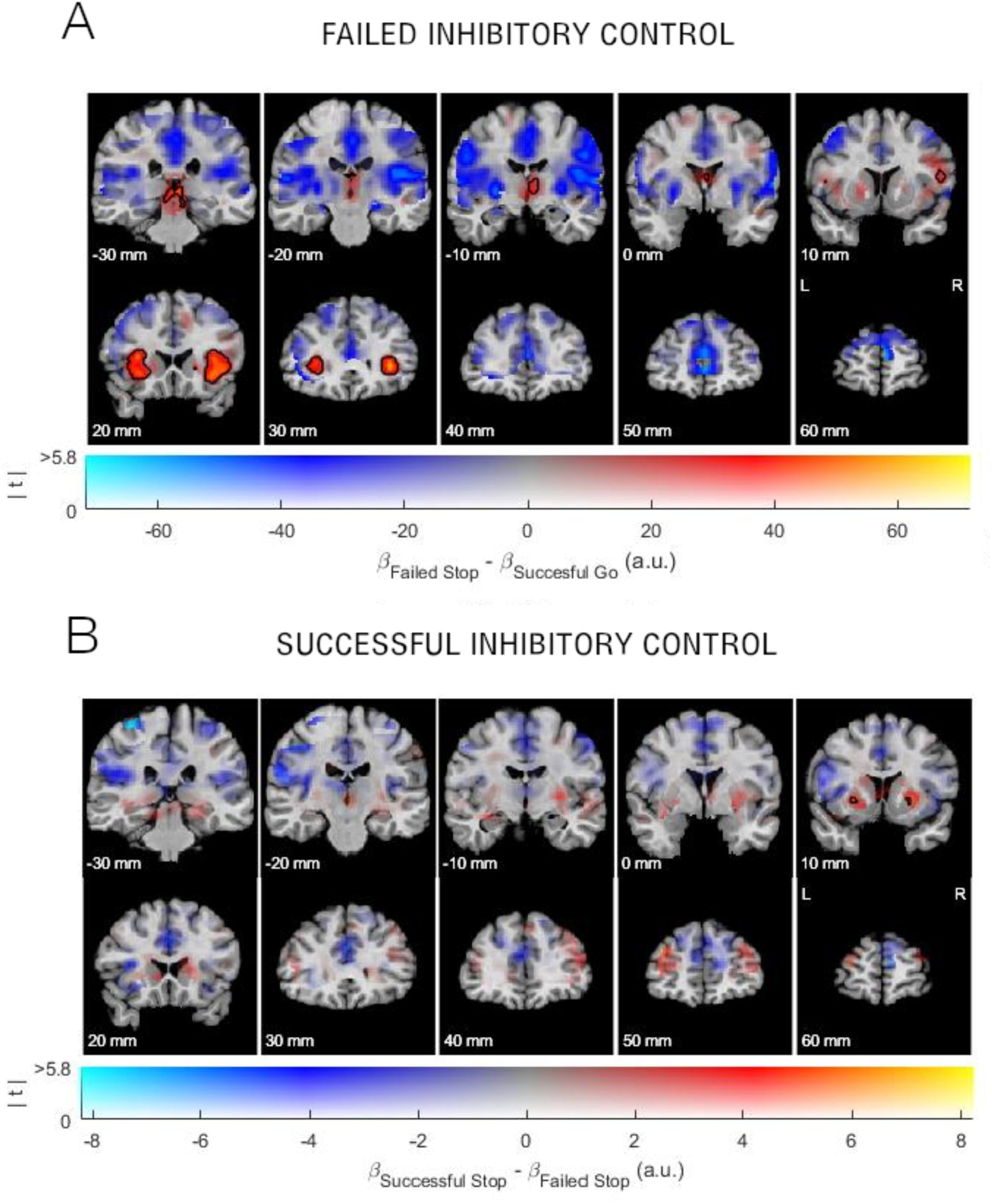
Task activation across all groups during (A) failed inhibitory control (failed stop – successful go) and (B) successful inhibitory control (failed – successful stop), which showed common patterns of activation. The colors reflect uncorrected activation, voxels outlined in black reflect survived correction at *p*_FWE_ = 0.05 showing fronto-striatal activation during cognitive control. The numbers above the color bars reflect t-values. Neuroimaging data are plotted using a procedure introduced by (56) and implemented by (57).

### Fronto-striatal functioning and glutamate

To investigate our main hypothesis regarding the association between fronto-striatal functioning and neurochemistry, we combined the ^1^H-MRS and fMRI data by extracting both beta values from our regions of interest as well as glutamate concentrations.

#### Failed inhibitory control

During failed inhibitory control, the interaction-term between striatal glutamate, diagnosis and time was associated with alterations in neural activation within the ACC (b= −4.89, t_(48.99)_ = −2.06, *p*= 0.044, *r* = 0.283). Compared to controls, the effect of striatal glutamate on neural activation in the ACC was decreased over time in the OCD group. However, when not considering striatal glutamate (two-way interaction between diagnosis and time), the OCD group showed an increase in neural activation compared to controls at T2 compared to T1 (b= 34.97, t_(48.66)_ = 2.07, *p*= 0.044, *r* = 0.285). While a two-way interaction term revealed significant results as well, the three-way interaction informs us that the interaction between striatal glutamate and diagnosis on neural activation within the ACC varied with time, as shown in Figure 5. No effects were found regarding the ASD group or the RBS-R scores (all *p*-values > 0.05). See Figure 5 for more details.

**Figure 5:**
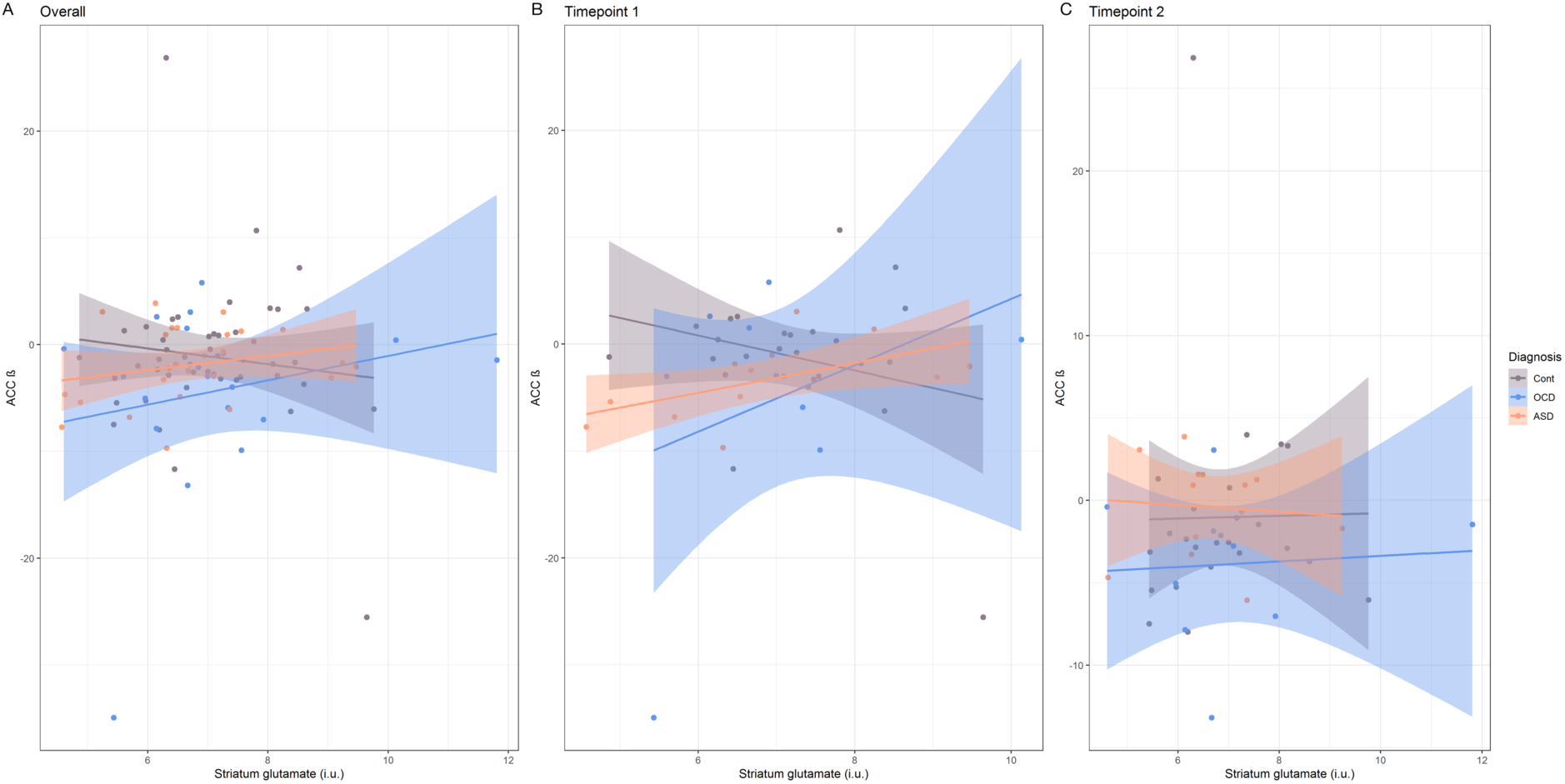
Effects of striatal glutamate on ACC BOLD signal during failed inhibitory control across timepoints (A), and separated over first time of measure (B) and second time of measure (C) during failed inhibitory control. The left plot shows the difference in the effects of striatal glutamate on ACC BOLD during failed inhibitory control in ASD (orange) and OCD (blue) compared to controls (grey). When separated over time of measure (B and C), the plots show the effects to be different in OCD compared to the control group over time. *Note:* This figure shows raw data-points, not model estimates.

#### Successful inhibitory control

During successful inhibitory control, there was a larger increase in striatal neural activity with an increase in ACC glutamate in the OCD group compared with controls (b = 0.19, t_(100)_ = 2.147, *p* = 0.036, *r* = 0.21), which was independent of time. The same positive association was found when including RBS-R compulsivity instead of diagnostic group ((b = 0.02, t_(102)_ = 2.394, *p* = 0.019, *r* = 0.23), indicating an increase in striatal activity with an increase in ACC glutamate when compulsivity scores are higher). No effects were found for the ASD group or with RBS-R total scores (all *p*-values > 0.05). A visual representation of these results can be seen in Figure 6.

**Figure 6:**
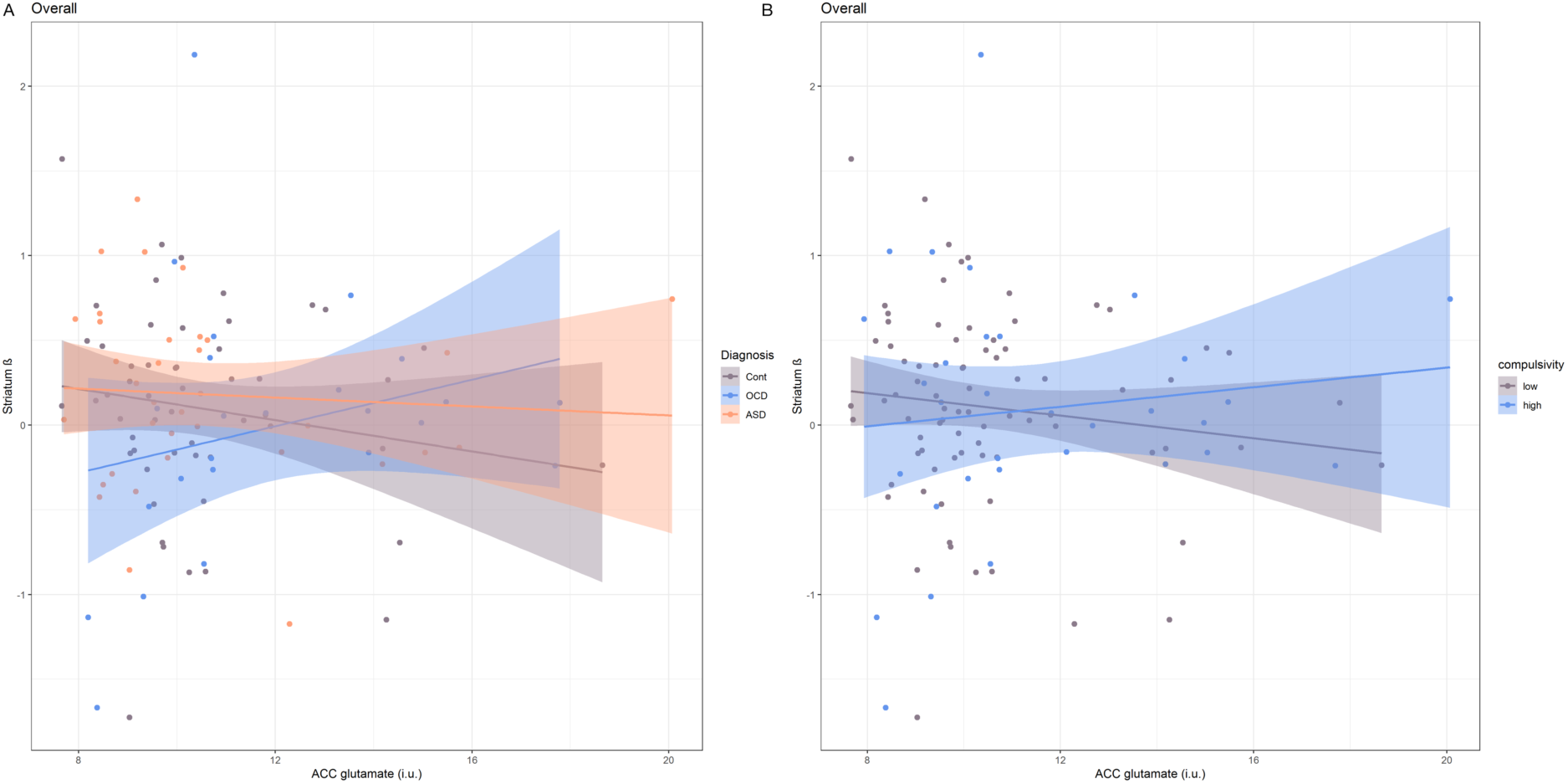
Effects of ACC glutamate on striatal BOLD signal during successful inhibitory control across timepoints combined. A: An increase in striatal activity during successful inhibitory control in the OCD group (blue) compared to the control group (grey) with an increase in ACC glutamate. B: Interaction between ACC glutamate and the continuous measure of RBS-R compulsivity (shown binary in high and low compulsivity, split by the overall mean score). *Note:* This figure shows raw data-points, not model estimates.

## Discussion

This is the first study that used a multi-center, longitudinal, transdiagnostic approach to investigate the associations of compulsivity with fronto-striatal glutamate concentrations and functioning during inhibitory control in a childhood/adolescent cross-disorder population.

Most interestingly, in our multi-modal analysis we found effects of fronto-striatal glutamate, diagnosis and time associated with alterations in fronto-striatal neural activity during both failed and successful inhibitory control. Neural activation in the ACC during failed inhibitory control was modulated by striatal glutamate, diagnosis and time. In the OCD group striatal glutamate showed a different, positive, modulatory effect compared to ASD and controls during T1, while at T2 this effect was diminished and resembled the pattern of the other groups. During successful inhibitory control, on the other hand, ACC glutamate was positively associated with striatal functioning in the OCD group versus controls, independent of time as well as with the continuous measure of compulsivity, but not the total score of the RBS-R. These results are partly in contrast with our previous study in which we showed an opposite relationship between ACC glutamate and striatal functioning (9). However, in the current study this association was related to measures of compulsivity. These results indicate that participants with OCD and/or high scores of compulsivity on a continuum seem to have lower neural activation in the striatum which reverses with the level of glutamate in the ACC. Considering the very limited research on these measures during adolescence, even more so in OCD than in ASD, these results are an important step towards increasing understanding of underlying mechanisms of development in compulsivity-related disorders.

We further found a larger decrease in ACC glutamate over time in ASD compared to controls. Previous studies investigating children with ASD have shown higher glutamate concentrations in ACC (49–51), while studies looking at adults with ASD have found both lower and higher glutamate concentrations in ACC compared to controls (7, 52). Our finding may therefore reflect changes in development into adulthood in ASD being different from development in controls. We found no such differences in the OCD group, although they did not significantly differ from the ASD group either, and previous studies with OCD have shown inconsistent results (7). This may be due to a larger heterogeneity in the disorder, and future studies considering possible subtypes of OCD may successfully disentangle such differing results. However, the previous study investigating an overlapping sample of our first wave measures (however, larger) at the first time-point (T1) found increased ACC glutamate in both ASD and OCD (8). Here we found no such differences over time, but we found effects of glutamate on neural activation during inhibitory control changing over time. This suggests there may also be differences in glutamate alterations during development between these disorders from childhood into adolescence. This, however, needs further investigation.

In the striatum we found that glutamate decreased over time in all groups. This is in line with the study that found no group differences in striatal glutamate during the first time of measure (8). Alterations in metabolite concentrations during development are known to occur in controls as well (53), and our finding may reflect such development in striatum, independent of a clinical diagnosis.

Our recent longitudinal TACTICS study on inhibitory control in ASD and OCD found improvements in SSRT over time, regardless of primary diagnosis (58). In our partly overlapping subsample in the current study, as shown in the supplementary material, males performed better than females. The fact that we did not find a general improvement may, however, be due to a lack of power and/or to a larger proportion of males in this subsample.

We found no whole-brain differences during inhibitory control in participants with ASD and OCD, which is also in concordance with the main findings from Gooskens and colleagues (58). However, other studies with similar behavioral results still found altered brain activation during inhibitory control (17, 25, 27).

Strengths of this study are combining categorical and dimensional analyses, as well as a longitudinal approach to investigate the relation between compulsivity with glutamate. Our study is also further strengthened by its multimodal nature, and investigating ASD and OCD together. There were also some limitations. Firstly, the OCD group was smaller than the ASD group, which may have led to less power and the possibility of false negatives. However, we still found significant associations with glutamate concentrations affecting functional activity in OCD. Secondly, the percentage GM in the striatum decreased over time, suggesting worse voxel placement. However, these changes were not different across diagnostic groups and therefore probably did not affect our main findings. We additionally had a large overall loss of sample size, due to lack of longitudinal data in several participants. There are also difficulties performing multicenter studies, where data quality may differ across sites. However, we did manage to control for these effects in our models and our results may not have been affected by left-over site effects.

The time between measures in this study was about one year. For future studies we suggest using an even longer time period between measures, extending into older ages, for an increased understanding of further development in ASD, OCD and compulsive behavior.

In conclusion, we found significant associations of increased glutamate concentrations in striatum, with decreased functional activity in ACC during failed inhibitory control in OCD, and that over time this effect changed differently in OCD compared to controls. We also found that ACC glutamate had a larger effect on striatal neural activation in participants with increased compulsive behavior (indicated by both the diagnostic and continuous finding) during successful inhibitory control. These results should be replicated in an independent sample, but this study has given new insights into the alterations of glutamate in ASD and OCD during development in adolescence, and its role in functional activity.

## Acknowledgements

The research leading to these results received funding from the European Community’s Seventh Framework Program (FP7/2007-2013) TACTICS under grant agreement no. 278948. This research was also supported by the Innovative Medicines Initiative Joint Undertaking under grant agreement number 115300 (EU-AIMS), resources of which are composed of financial contribution from the European Union’s Seventh Framework Programme (FP7- /2007 - 2013) and the European Federation of Pharmaceutical Industries and Associations (EFPIA) companies’ in kind contribution. J Naaijen is supported by a VENI grant of the Netherlands Organization for Scientific Research (NWO, grant number VI.Veni.194.032). The authors would like to thank Nicole Driessen, Saskia de Ruiter, Sophie Akkermans, Vincent Mensen, Muriel Bruchhage, Isabella Wolf and Regina Boecker-Schlier for their help in data-collection and all participants for their participation.

## References

1. American Psychiatric Association (2013): Diagnostic and statistical manual of mental disorders (5th ed). Washington, DC:Author.. doi: 10.1176/appi.books.9780890425596.744053.

2. Robbins TW, Gillan CM, Smith DG, de Wit S, Ersche KD (2012): Neurocognitive endophenotypes of impulsivity and compulsivity: Towards dimensional psychiatry. Trends Cogn Sci. 16: 81–91.

3. Anholt GE, Cath DC, Van Oppen P, Eikelenboom M, Smit JH, Van Megen H, Van Balkom AJLM (2010): Autism and adhd symptoms in patients with ocd: Are they associated with specific oc symptom dimensions or oc symptom severity. J Autism Dev Disord.. doi: 10.1007/s10803-009-0922-1.

4. Zandt F, Prior M, Kyrios M (2007): Repetitive behaviour in children with high functioning autism and obsessive compulsive disorder. J Autism Dev Disord.. doi: 10.1007/s10803-006-0158-2.

5. Chamberlain SR, Menzies L (2009): Endophenotypes of obsessive-compulsive disorder: Rationale, evidence and future potential. Expert Rev Neurother.. doi: 10.1586/ern.09.36.

6. Chamberlain SR, Fineberg NA, Menzies LA, Blackwell AD, Bullmore ET, Robbins TW, Sahakian BJ (2007): Impaired cognitive flexibility and motor inhibition in unaffected first-degree relatives of patients with obsessive-compulsive disorder. Am J Psychiatry.. doi: 10.1176/ajp.2007.164.2.335.

7. Naaijen J, Lythgoe DJ, Amiri H, Buitelaar JK, Glennon JC (2015): Fronto-striatal glutamatergic compounds in compulsive and impulsive syndromes: A review of magnetic resonance spectroscopy studies. Neurosci Biobehav Rev. 52.

8. Naaijen J, Zwiers MP, Amiri H, Williams SC, Durston S, Oranje B, et al. (2016): Fronto-Striatal Glutamate in Autism Spectrum Disorder and Obsessive Compulsive Disorder. Neuropsychopharmacology. 1–35.

9. Naaijen J, Lythgoe DJ, Zwiers MP, Hartman CA, Hoekstra PJ, Buitelaar JK, Aarts E (2018): Anterior cingulate cortex glutamate and its association with striatal functioning during cognitive control. Eur Neuropsychopharmacol. 28. doi: 10.1016/j.euroneuro.2018.01.002.

10. Bari A, Robbins TW (2013): Inhibition and impulsivity: Behavioral and neural basis of response control. Prog Neurobiol.. doi: 10.1016/j.pneurobio.2013.06.005.

11. Chmielewski WX, Beste C (2015): Action control processes in autism spectrum disorder – Insights from a neurobiological and neuroanatomical perspective. Prog Neurobiol.. doi: 10.1016/j.pneurobio.2014.11.002.

12. Fineberg N a, Potenza MN, Chamberlain SR, Berlin H a, Menzies L, Bechara A, et al. (2010): Probing compulsive and impulsive behaviors, from animal models to endophenotypes: a narrative review. Neuropsychopharmacology. 35: 591–604.

13. Aron AR, Durston S, Eagle DM, Logan GD, Stinear CM, Stuphorn V (2007): Converging evidence for a fronto-basal-ganglia network for inhibitory control of action and cognition. J Neurosci.. doi: 10.1523/JNEUROSCI.3644-07.2007.

14. Aron AR, Poldrack RA (2006): Cortical and subcortical contributions to stop signal response inhibition: Role of the subthalamic nucleus. J Neurosci.. doi: 10.1523/JNEUROSCI.4682-05.2006.

15. Dalley JW, Everitt BJ, Robbins TW (2011): Impulsivity, Compulsivity, and Top-Down Cognitive Control. Neuron.. doi: 10.1016/j.neuron.2011.01.020.

16. Zandbelt BB, Vink M (2010): On the role of the striatum in response inhibition. PLoS One.. doi: 10.1371/journal.pone.0013848.

17. Chantiluke K, Barrett N, Giampietro V, Santosh P, Brammer M, Simmons A, et al. (2015): Inverse fluoxetine effects on inhibitory brain activation in non-comorbid boys with ADHD and with ASD. Psychopharmacology (Berl).. doi: 10.1007/s00213-014-3837-2.

18. Albajara Sáenz A, Septier M, Van Schuerbeek P, Baijot S, Deconinck N, Defresne P, et al. (2020): ADHD and ASD: distinct brain patterns of inhibition-related activation? Transl Psychiatry. 10. doi: 10.1038/s41398-020-0707-z.

19. Gooskens B, Bos DJ, Mensen V, Shook D, Bruchhage M, Naaijen J, et al. (2018): No evidence of differences in cognitive control in children with autism spectrum disorder or obsessive-compulsive disorder: An fMRI study..

20. Verbruggen F, Logan GD (2008): Response inhibition in the stop-signal paradigm. Trends Cogn Sci.. doi: 10.1016/j.tics.2008.07.005.

21. De Wit SJ, De Vries FE, Van Der Werf YD, Cath DC, Heslenfeld DJ, Veltman EM, et al. (2012): Presupplementary motor area hyperactivity during response inhibition: A candidate endophenotype of obsessive-compulsive disorder. Am J Psychiatry.. doi: 10.1176/appi.ajp.2012.12010073.

22. Penadés R, Catalán R, Rubia K, Andrés S, Salamero M, Gastó C (2007): Impaired response inhibition in obsessive compulsive disorder. Eur Psychiatry.. doi: 10.1016/j.eurpsy.2006.05.001.

23. Marzuki AA, Pereira de Souza AMFL, Sahakian BJ, Robbins TW (2020): Are candidate neurocognitive endophenotypes of OCD present in paediatric patients? A systematic review. Neurosci Biobehav Rev.. doi: 10.1016/j.neubiorev.2019.12.010.

24. Apergis-Schoute AM, Bijleveld B, Gillan CM, Fineberg NA, Sahakian BJ, Robbins TW (2018): Hyperconnectivity of the ventromedial prefrontal cortex in obsessive-compulsive disorder. Brain Neurosci Adv.. doi: 10.1177/2398212818808710.

25. Woolley J, Heyman I, Brammer M, Frampton I, McGuire PK, Rubia K (2008): Brain activation in paediatric obsessive-compulsive disorder during tasks of inhibitory control. Br J Psychiatry.. doi: 10.1192/bjp.bp.107.036558.

26. Roth RM, Saykin AJ, Flashman LA, Pixley HS, West JD, Mamourian AC (2007): Event-Related Functional Magnetic Resonance Imaging of Response Inhibition in Obsessive-Compulsive Disorder. Biol Psychiatry.. doi: 10.1016/j.biopsych.2006.12.007.

27. Rubia K, Cubillo A, Smith AB, Woolley J, Heyman I, Brammer MJ (2010): Disorder-specific dysfunction in right inferior prefrontal cortex during two inhibition tasks in boys with attention-deficit hyperactivity disorder compared to boys with obsessive-compulsive disorder. Hum Brain Mapp.. doi: 10.1002/hbm.20864.

28. Silverman JM, Buxbaum JD, Ramoz N, Schmeidler J, Reichenberg A, Hollander E, et al. (2008): Autism-related routines and rituals associated with a mitochondrial aspartate/glutamate carrier SLC25A12 polymorphism. Am J Med Genet Part B Neuropsychiatr Genet.. doi: 10.1002/ajmg.b.30614.

29. Naaijen J, de Ruiter S, Zwiers MP, Glennon JC, Durston S, Lythgoe DJ, et al. (2016): COMPULS: Design of a multicenter phenotypic, cognitive, genetic, and magnetic resonance imaging study in children with compulsive syndromes. BMC Psychiatry. 16. doi: 10.1186/s12888-016-1072-6.

30. American Psychiatric Association (2000): Diagnostic and statistical manual of mental disorders: DSM-IV-TR (text revision). Am J Psychiatry..

31. Lord C, Rutter M, Le Couteur A (1994): Autism Diagnostic Interview-Revised: A revised version of a diagnostic interview for caregivers of individuals with possible pervasive developmental disorders. J Autism Dev Disord.. doi: 10.1007/BF02172145.

32. Scahill L, Riddle MA, McSwiggin-Hardin M, Ort SI, King RA, Goodman WK, et al. (1997): Children’s Yale-Brown Obsessive Compulsive Scale: Reliability and validity. J Am Acad Child Adolesc Psychiatry.. doi: 10.1097/00004583-199706000-00023.

33. Bordin I a, Rocha MM, Paula CS, Teixeira MCT V, Achenbach TM, Rescorla L a, Silvares EFM (2013): Child Behavior Checklist (CBCL),Youth Self-Report (YSR) and Teacher’s Report Form(TRF): an overview of the development of the original and Brazilian versions. Cad Saude Publica. 29: 13–28.

34. Lam KSL, Aman MG (2007): The repetitive behavior scale-revised: Independent validation in individuals with autism spectrum disorders. J Autism Dev Disord.. doi: 10.1007/s10803-006-0213-z.

35. Rubia K, Smith AB, Brammer MJ, Taylor E (2003): Right inferior prefrontal cortex mediates response inhibition while mesial prefrontal cortex is responsible for error detection. Neuroimage.. doi: 10.1016/S1053-8119(03)00275-1.

36. Jack CR, Bernstein MA, Fox NC, Thompson P, Alexander G, Harvey D, et al. (2008): The Alzheimer’s Disease Neuroimaging Initiative (ADNI): MRI methods. J Magn Reson Imaging.. doi: 10.1002/jmri.21049.

37. Jack CR, Bernstein MA, Borowski BJ, Gunter JL, Fox NC, Thompson PM, et al. (2010): Update on the Magnetic Resonance Imaging core of the Alzheimer’s Disease Neuroimaging Initiative. Alzheimer’s Dement.. doi: 10.1016/j.jalz.2010.03.004.

38. Haase A, Frahm J, Hanicke W, Matthaei D (1985): 1H NMR chemical shift selective (CHESS) imaging. Plasma Sources Sci Technol.. doi: 10.1088/0031-9155/30/4/008.

39. Jenkinson M, Bannister P, Brady M, Smith S (2002): Improved optimization for the robust and accurate linear registration and motion correction of brain images. Neuroimage.. doi: 10.1016/S1053-8119(02)91132-8.

40. Pruim RHR, Mennes M, van Rooij D, Llera A, Buitelaar JK, Beckmann CF (2015): ICA-AROMA: A robust ICA-based strategy for removing motion artifacts from fMRI data. Neuroimage. 112: 267–277.

41. Pruim RHR, Mennes M, Buitelaar JK, Beckmann CF (2015): Evaluation of ICA-AROMA and alternative strategies for motion artifact removal in resting state fMRI. Neuroimage. 112: 278–287.

42. Greve DN, Fischl B (2009): Accurate and robust brain image alignment using boundary-based registration. Neuroimage.. doi: 10.1016/j.neuroimage.2009.06.060.

43. Andersson JLR, Jenkinson M, Smith S (2007): Non-linear registration aka spatial normalisation. FMRIB Tech Rep TRO7JA2..

44. Provencher S (2014): Manual..

45. Provencher SW (2001): Automatic quantitation of localized in vivo 1H spectra with LCModel. NMR Biomed. 14: 260–264.

46. Booth DS, Szmidt-Middleton H, King N, Westbrook MJ, Young SL, Kuo A, et al. (2018): RStudio: Integrated Development for R. Nature.. doi: 10.1108/eb003648.

47. Bates D, Mächler M, Bolker BM, Walker SC (2015): Fitting linear mixed-effects models using lme4. J Stat Softw.. doi: 10.18637/jss.v067.i01.

48. Kreis R (2016): The trouble with quality filtering based on relative Cramér-Rao lower bounds. Magn Reson Med. 75: 15–18.

49. Bejjani A, O’Neill J, Kim J a, Frew AJ, Yee VW, Ly R, et al. (2012): Elevated glutamatergic compounds in pregenual anterior cingulate in pediatric autism spectrum disorder demonstrated by 1H MRS and 1H MRSI. PLoS One. 7: e38786.

50. Hassan TH, Abdelrahman HM, Abdel NR, El-masry NM, Hashim HM, El-gerby KM, Abdel NR (2013): Research in Autism Spectrum Disorders Blood and brain glutamate levels in children with autistic disorder. Res Autism Spectr Disord. 7: 541–548.

51. Joshi G, Biederman J, Wozniak J, Goldin RL, Crowley D, Furtak S (2012): Magnetic resonance spectroscopy study of the glutamatergic system in adolescent males with high-functioning autistic disorder: a pilot study at 4T.. doi: 10.1007/s00406-012-0369-9.

52. Ford T, Crewther D (2016): A comprehensive review of the 1H-MRS metabolite spectrum in autism spectrum disorder. Front Mol Neurosci. 9.

53. Horská A, Kaufmann WE, Brant LJ, Naidu S, Harris JC, Barker PB (2002): In vivo quantitative proton MRSI study of brain development from childhood to adolescence. J Magn Reson Imaging.. doi: 10.1002/jmri.10057.

54. Ginestet C (2011): ggplot2: Elegant Graphics for Data Analysis. J R Stat Soc Ser A (Statistics Soc.. doi: 10.1111/j.1467-985x.2010.00676_9.x.

55. Hintze JL, Nelson RD (1998): Violin plots: A box plot-density trace synergism. Am Stat.. doi: 10.1080/00031305.1998.10480559.

56. Allen EA, Erhardt EB, Calhoun VD (2012): NeuroView Data Visualization in the Neurosciences: Overcoming the Curse of Dimensionality NeuroView. Neuron. 74: 603–608.

57. B Z (2017): Slice Display figure. Figshare. 106084/m9.figshare4742866..

58. Gooskens B, Bos DJ, Naaijen J, Akkermans SEA, Kaiser A, Hohmann S, Bruchhage NMK, Banachewski T, Brandeis D, Williamsn SCR, Lythgoe DJ, Buitelaar JK, Oranje B, Durston S, The TACTICS consotrium (2020). The development of cognitive control in children with Autsim spectrum disorders or obsessive-compulsive disorder: A longitudinal fMRI study. BioRxiv. doi: https://doi.org/10.1101/2020.04.09.033696

