## Supplementary material for "Longitudinal alterations in fronto-striatal glutamate are associated with functioning during inhibitory control in autism spectrum disorder and obsessive compulsive disorder"

**Supplementary information**

Multimodal imaging of compulsivity across neurodevelopmental disorders: A longitudinal investigation

Hollestein et al.

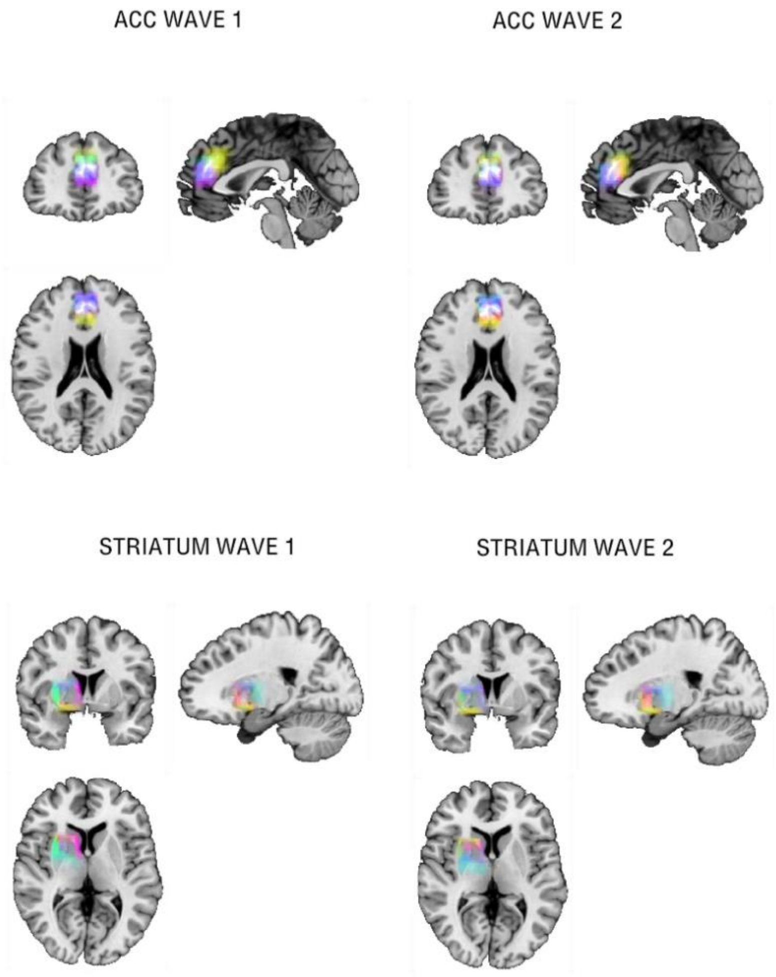


**Figure S1:** Superposition on the MNI152 template of all individual voxel placements in ACC and striatum, for all sites (London, blue; Mannheim, yellow; Nijmegen, pink). The placements are consistent across and within sites.

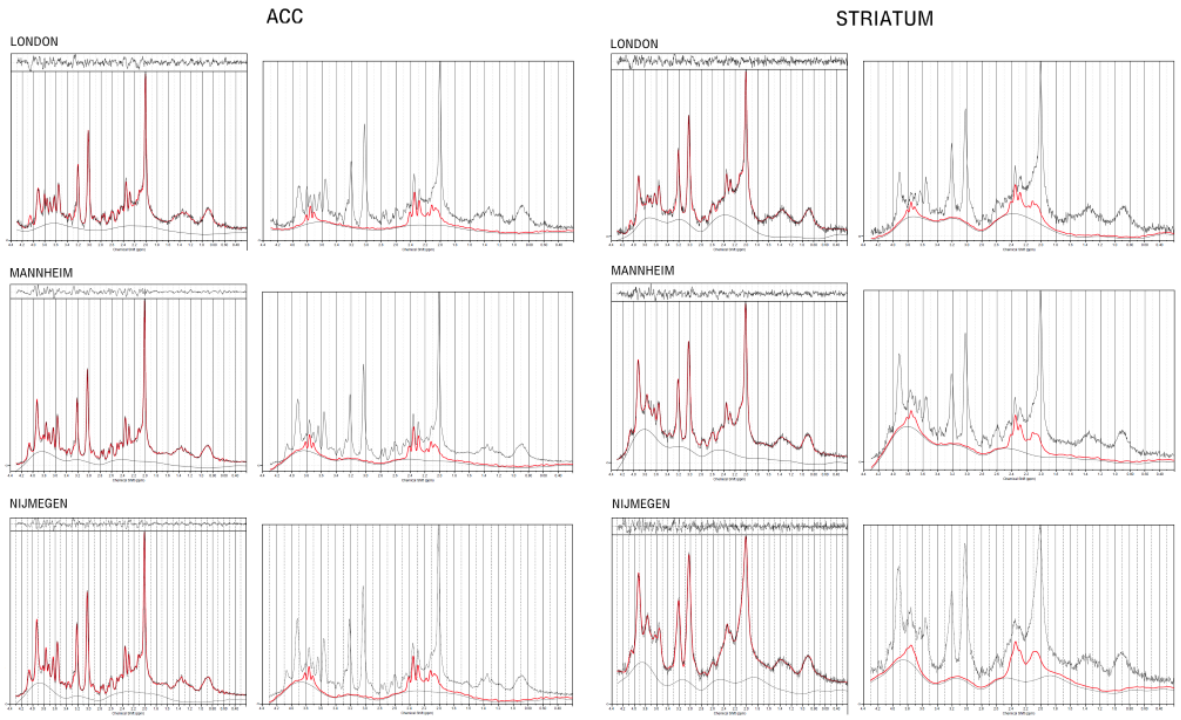


**Figure S2:** Example spectra of a 3T from proton magnetic resonance spectroscopy (1H-MRS) Linear Combination (LC) Model spectral fit in ACC and striatum across all sites. The top of the images represents the residuals. The black line represents frequency-domain data, the red line is the LCModel fit. The right images show the fits for glutamate only.

**Details main analysis**

Linear mixed effects models were used for our statistical analyses using the lme4 package (Bates et al., 2014) in R (RStudio Team, 2016). The lmer function was used to fit linear mixed-effects models. For analysis of associations of glutamate concentrations in ACC and striatum with BOLD signal in the same areas, the same model was used for all contrasts of interest:

$$\text{BOLD } \beta_1 \sim \text{Glutamate}_2 * \text{Diagnosis} + \text{Site} + (1 \mid \text{Participant})$$

[1,2]The same model was used for both ACC and striatum

For analysis of glutamate concentrations in ACC and striatum associated with diagnosis or compulsivity, these codes were used:

$$\text{Glutamate concentration}_1 \sim \text{Diagnosis} * \text{Time} + \text{Site} + (1 \mid \text{Participant})$$

$$\text{Glutamate concentration}_1 \sim \text{RBS-Total score} * \text{Time} + \text{Site} + (1 \mid \text{Participant})$$

$$\text{Glutamate concentration}_1 \sim \text{RBS-Compulsivity} * \text{Time} + \text{Site} + (1 \mid \text{Participant})$$

[1]The same model was used for both ACC and striatum

For analysis of SSRT group comparison, over time this code was used:

$$\text{SSRT} \sim \text{Diagnosis} * \text{Time} + \text{Site} + (1 \mid \text{Participant})$$

**Medication use over time**

During the first time of measure, in the ASD group two people used stimulants, and one anti-depressants. In the OCD group five people used antidepressants and one anti-psychotics. In the second time of measure, one of the participants using stimulants and the participant using anti-depressants now also used antipsychotics. An additional participant used antidepressants, and one antipsychotics and stimulants. In the OCD group two were no longer on antidepressants, the one using antipsychotics in the first time of measure now also used antidepressants, and one participant had started using stimulants. None of the controls used medication at any time of measure.

**Stop-Signal task (behavioral)**

**Analysis.** The behavioral measure of interest on the SST was the stop-signal reaction time (SSRT), which was calculated using the integration method (1, 2), where the reaction time (RT) of correct go trials was rank ordered, then the *n*th go-RT was selected, where *n* was derived by multiplying the number of correct go-trials by the probability that the participant respond to a stop signal. The SSRT was then estimated by subtracting the mean SSD from the *n*th go-RT (3). Participants were excluded from analysis for excessive motion or when they showed an SSRT < 50 ms as it is indicative of not performing the task properly, for example by constantly pressing buttons without paying attention to cues which results in atypically short response times on correct go-trials. This resulted in 41 participants included for stop-task analysis (ASD = 12, OCD = 8, controls = 21). Data from T1 and T2 were initially analyzed separately allowing investigation of group differences without the possible influence of time. Shapiro-Wilk normality tests showed that there was no normal distribution in neither time of measure, and therefore Fligner-Killeen tests of homogeneity of variance were used, which showed that there was equal variance between diagnosis groups in both T1 and T2. Consequently, Kruskal-Wallis tests were used to analyze differences in SSRT between groups (ASD, OCD or controls) for T1 and T2 independently. To investigate behavioral differences between groups on the stop-task measured by the SSRT, Kruskal-Wallis tests were used to compare groups, and a mixed effects model was used to analyze changes between groups over time of measure, including the same covariates as described before.

**Results.** There were no group differences in SSRT in T1 ($c_{2(2)}$= 2.84, $p$ > 0.1) or T2($c_{2(2)}$=2.64, $p$ > 0.1), showing similar performance across groups. Across T1 and T2 a significant effect of sex was found (b = -101.34, $t_{(34.3)}$=-2.21 $p$ = 0.03, $r$ = 0.35), indicating an better stop-task performance in males, compared to females.

**Supplemental references**


1. Verbruggen F, Aron AR, Band GPH, Beste C, Bissett PG, Brockett AT, *et al.* (2019): A consensus guide to capturing the ability to inhibit actions and impulsive behaviors in the stop-signal task. *Elife*. . doi: 10.7554/eLife.46323.

2. Verbruggen F, Chambers CD, Logan GD (2013): Fictitious Inhibitory Differences: How Skewness and Slowing Distort the Estimation of Stopping Latencies. *Psychol Sci*. . doi: 10.1177/0956797612457390.

3. Verbruggen F, Logan GD (2008): Response inhibition in the stop-signal paradigm. *Trends Cogn Sci*. . doi: 10.1016/j.tics.2008.07.005.
